# Treatment with Tumor-Treating Fields (TTFields) Suppresses Intercellular Tunneling Nanotube Formation *In Vitro* and Upregulates Immuno-Oncologic Biomarkers *In Vivo* in Malignant Mesothelioma

**DOI:** 10.1101/2022.12.29.522223

**Authors:** Akshat Sarkari, Sophie Korenfeld, Karina Deniz, Katherine Ladner, Phillip Wong, Sanyukta Padmanabhan, Rachel I Vogel, Laura Sherer, Naomi Courtemanche, Clifford J Steer, Kerem Wainer-Katsir, Emil Lou

## Abstract

Intercellular communication is critical for the development of invasive cancers. Multiple forms of intercellular communication have been well characterized, involving diffusible soluble factors or contact-dependent channels for immediately adjacent cells. Over the past 1-2 decades, the emergence of a unique form of F-actin-based cellular protrusion known as tunneling nanotubes (TNTs) has filled the niche of long-range cell-contact dependent intercellular communication that facilitates cell growth, differentiation, and in the case of invasive cancer phenotypes, a more chemoresistant phenotype. The cellular machinery of TNT-mediated transport is an area of active investigation, and microtubules have been implicated in this process as they are in other membranous protrusions. Tumor-Treating Fields (TTFields) therapy is a novel therapeutic strategy in clinical use for patients with advanced cancers, based on the principle of using low-intensity alternating electric fields to disrupt microtubules in cancer cells undergoing mitosis. Other mechanisms of action have also been demonstrated. In this study, we investigated the effects of TTFields on TNTs in malignant pleural mesothelioma (MPM) *in vitro* and also on the spatial transcriptomic landscape *in vivo*. We found that applying TTFields at 1.0 V/cm significantly suppressed TNT formation in a biphasic MPM cell line (MSTO-211H), but not in sarcomatoid MPM (VAMT). At these parameters, TTFields significantly reduced cell count in MSTO-211H, but did not significantly alter intercellular transport of mitochondria via intact TNTs. To understand how TTFields may impact expression of genes with known involvement to TNT formation and overall tumor growth, we performed spatial genomic assessment of TTFields-treated tumors from an *in vivo* animal model of MPM, and detected upregulation of immuno-oncologic biomarkers with simultaneous downregulation of pathways associated with cell hyperproliferation, invasion, and other critical regulators of oncogenic growth. Several molecular classes and pathways coincide with markers that we and others have found to be differentially expressed in cancer cell TNTs, including MPM specifically. In this study, we report novel cellular and molecular effects of TTFields in relation to tumor communication networks enabled by TNTs and related molecular pathways. These results position TNTs as potential therapeutic targets for TTFields-directed cancer treatment strategies; and also identify the ability of TTFields to potentially remodel the tumor microenvironment, thus enhancing response to immunotherapeutic drugs.

## Introduction

Intercellular communication in the dense and highly heterogeneous tumor matrix is a critical function and hallmark of invasive cancers. Multiple forms of intercellular communication have been well documented and characterized, including gap junctions, extracellular vesicles, and signaling via diffusible growth factors, among others. In the past decade, a unique form of F-actin-based cellular protrusion known as tunneling nanotubes (TNTs) has been shown to mediate direct contact-dependent intercellular communication in many cell types, and particularly, invasive cancer phenotypes. The cellular machinery of TNT-mediated transport is an area of active investigation, and microtubules have been implicated in this process as they are in other membranous protrusion.

Tunneling nanotubes (TNTs) are F-actin-based membrane protrusions that physically connect cells over distances that are typically 100-500 µm or longer^1, 2, 3, 4, 5, 6^. These ultrafine structures were first characterized in 2004 in PC12 cells, a cell line derived from rat pheochromocytoma^7^, and are morphologically and functionally distinct from other membranous protrusions such as filopodia or lamellipodia, which aid in motility and attachment to the extracellular matrix (ECM)^8^. Unlike filopodia and lamellipodia, TNTs are non-adherent to the substratum in cells cultured *in vitro*^6, 7, 9, 10^. TNTs have been identified in many forms of cells, including fibroblasts, epithelial cells, and neurons, but are prominently upregulated in cancer cells^2, 6, 9, 11, 12, 13, 14^. The potential for a single TNT to remain stable for hours, combined with its upregulation in cancer phenotypes, indicates that TNTs may be capable of mounting a rapid communication response to external stimuli or insults, including chemotherapeutic drugs^6, 9^. However, the mechanism(s) of TNT formation and the role of actin in TNT formation and stability across cell types remain largely unknown.

Tumor-Treating Fields (TTFields) therapy is a novel therapeutic strategy in clinical use for patients with several forms of advanced cancers, including glioblastomas and malignant pleural mesotheliomas (MPM). It is based on the principle of using low-intensity alternating electric fields to disrupt microtubules in cancer cells undergoing mitosis. These fields apply forces on charges and polarizable molecules inside and around cells.

TTFields can disrupt mitosis in malignant cells due to its ability to interfere with mitotic spindle assembly through impairment of microtubule polymerization^15^. Microtubules are essential in ensuring that chromosomes attach and segregate correctly during metaphase and anaphase, respectively. The individual subunits of microtubules, known as tubulins, are heterodimers with 2 distinct protein domains, in which one has a positive end and one has a negative end, creating a dipole^16^. If microtubules are not allowed to polymerize, cell division cannot occur, and this is typically followed by chromosomal abnormalities and mitotic cell death^17^.

Additionally, TTFields application creates a nonuniform electric field during the telophase phase of mitosis due to alignment of the cell in cytokinesis, leading to a process known as dielectrophoresis, which can also result in improper cell division and mitotic death^18^. Other mechanisms of action have also been demonstrated, including downregulation of DNA damage response, impairment of cancer cell migration, and induction of anti-cancer immunity^19, 20^. Unlike systemic cancer therapies, TTFields delivery is focused on the tumor area, thus minimizing effects on non-malignant cells outside the treated area. Due to differences in geometrical and electrical properties, the TTFields frequency range is deleterious to cancer but not to benign cells and is optimized to a specific tumor type^18, 20^. This technology is currently applied concomitantly with standard-of-care treatment approaches for patients with glioblastoma and mesothelioma, with clinical trials also ongoing in many other forms of metastatic or difficult-to-treat forms of cancer.

The bulk of studies to date on cellular effects and mechanism(s) of TTFields has focused on disruption of microtubules, leading to decreased cell division. For this study we hypothesized that TTFields also affects formation of F-actin based TNTs in intact cells. We have previously reported that TNTs are significantly upregulated in multiple forms of MPM, which serves as an excellent model system for studying and characterizing cellular structure, function, and dynamics of TNT-mediated intercellular communication of cellular signals. Here, we report the effects of TTFields on TNTs connecting MPM cells *in vitro*, and on cell-free monomeric and filamentous actin. We also examined differential expression of gene pathways of immune response, proliferative growth, and other hallmarks of MPM in an *in vivo* animal model in order to elucidate the impact of TTFields on the expression of genes known to be involved in TNT formation.

## Results

### Establishing TTFields application impact on TNT formation in malignant mesothelioma cells

We utilized two mesothelioma cell lines, MSTO-211H and VAMT, to investigate effects of TTFields application on TNT formation and function. These two cell lines were used having previously demonstrated that they reliably and reproducibly form TNTs in culture under variable conditions, and are thus ideal for *in vitro* studies. TTFields were applied to cells *in vitro* using two devices: inovitro™, which applies TTFields to cells in culture to which the electrodes from the power supply provide a pre-specified level of intensity and frequency of the alternating electric fields; and inovitro Live™, in which the configuration is adapted for continuous administration of TTFields while permitting time-lapse microscopic imaging. We first tested the inovitro Live device to treat MSTO-211H at differing frequencies to establish parameters used to impact TNT formation.

Previously, bidirectional application of TTFields has shown increased cytotoxicity relative to unidirectional delivery^21^, with highest cytotoxicity for MSTO-211H cells displayed at a frequency of 150 kHz^22^. Instead, we sought to elucidate the initial impact of TTFields on TNT protrusion formation, which may require differing frequencies than what is demonstrated to be most effective for a cytotoxic effect. Thus, we tested differing frequencies and directional vectors for TTFields application. TTFields intensity was administered at 1.0 V/cm but a frequency of either 200 kHz or 150 kHz was delivered bidirectionally or unidirectionally over a 72-hour period to MSTO-211H cells; these two frequencies were selected for testing because the approved devices for TTFields therapeutic delivery is applied at these frequencies (Fig 1A). We found that by 24 hours, unidirectional TTFields treatment at 200 kHz had fewer TNTs than the control (p = 0.004) and bidirectional application at 150 kHz (p = 0.005). As compared to control, bidirectional application at 200 kHz also had statistically significantly fewer TNTs (p < 0.0001). At times 48 and 72 hours, we also observed the decline in TNTs. As with previous studies, once cells become densely packed, they form fewer TNTs^23^. Together the data indicated that applying TTFields at 200 kHz unidirectionally is most effective at decreasing TNT formation in MSTO-211H cells and we utilized this frequency for the rest of our experiments.

**Figure 1.**
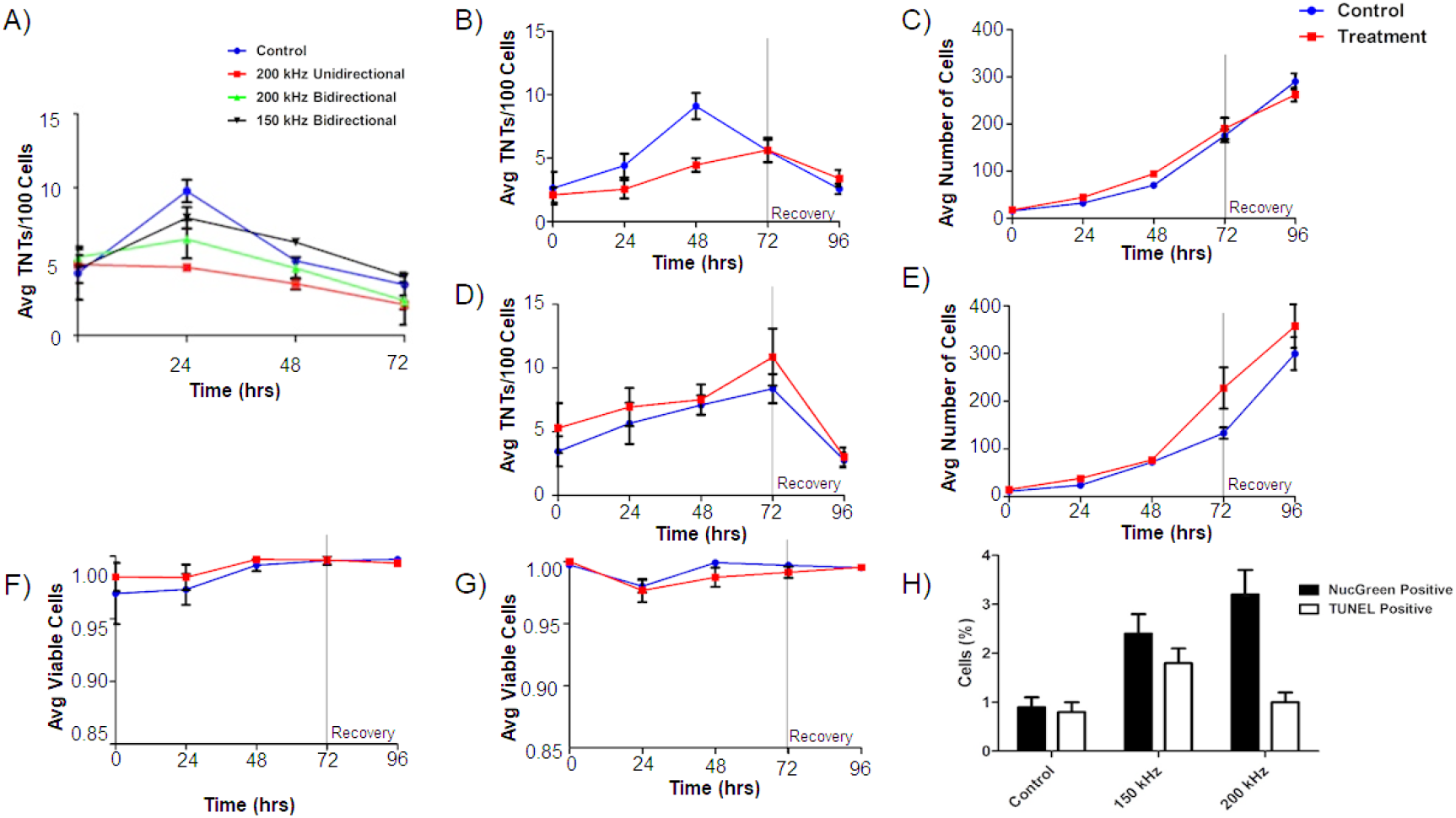
TNT formation, cell growth, and cell viability in MSTO-211H and VAMT. A) TNT formation in MSTO-211H following continuous TTFields exposure at 1.0 V/cm while varying frequency and field direction. 40,000 MSTO-211H cells were plated in a 35 mm dish and exposed to TTFields treatment at 1.0 V/cm with the above varying parameters; media was changed every 24 hours. B-C) TNT formation and cell growth in MSTO-211H following TTFields exposure when compared to control. As above, 40,000 cells were plated and were exposed continuously to TTFields bidirectionally; at 72 hours, TTFields treatment was discontinued to assess recovery of TNT formation (n=3). D-E) TNT formation and cell growth in VAMT following TTFields exposure with methodology as listed in B-C (n=3). F-G) Cell viability in both MSTO-211H (F) and VAMT (G) respectively following TTFields exposure. Cell viability and cytotoxicity was measured through NucGreen Dead 488 expression, which assesses loss of plasma membrane integrity. 7 random fields of view were selected and the ratio of live:dead cells was recorded (n=3). H) Cell viability measured by TUNEL assay and NucGreen Dead 488 expression in MSTO-211H exposed to TTFields at 150 and 200 kHz. MSTO-211H cells were treated with TTFields for 48 hours at either 150 kHz or 200 kHz. At the 48 hr timepoint, cell viability was measured through the TUNEL assay or through measuring fluorescent expression of Nuc Green Dead 488. The percentage of nonviable cells was graphed as compared to a control. A representative image of TUNEL positive control is displayed in Supplemental Figure 2. Cell count data are shown in Supplemental Figure 3. Statistical significance was assessed as a result of three independent experiments, with a linear mixed model used in Fig 1A and heteroscedastic t-test used in B.

### Effects of TTFields application on TNT formation in malignant mesothelioma cells

Next, we tested the inovitro device to treat MSTO-211H and VAMT cells independently plated on treated coverslips using a low intensity of 0.5 V/cm TTFields treatment over a 72-hour period. TTFields did not significantly alter the number of TNTs or cells at this intensity in either cell line (Fig S1). Once our experimental set-up was calibrated, we assessed effects of TTFields applied at an intensity of 1.0 V/cm with a 200 kHz frequency bidirectionally to evaluate the potential impact on TNT formation. Although we demonstrated the highest impact on TNT formation with unidirectional fields, we desired to emulate clinical conditions and efficacy as closely as possible and thus utilized bidirectional electric fields. Both cell lines were treated with TTFields over a 72-hour period to assess TNT formation and cell growth, with further assessment for an additional 24 hours after TTFields was discontinued to observe any latent effect or recovery of TNT formation. Over the 72-hour treatment period, we noted a statistically significant difference in TNT formation at 48 hours with MSTO-211H cells, but this difference was not present at 72 hours (Fig 1B, p=0.018).

Additionally, over the 24 hours following treatment stoppage, TNT formation decreased further in both the control and treatment groups and cell density continued to increase (Fig 1B,C). In fact, cell growth increased steadily in both treatment and control groups at nearly exponential rates, to reach confluency by the end of the experiments, indicating there was no latent effect on either TNT formation or cell growth from TTFields application. Unlike MSTO-211H, when VAMT cells were subjected to TTFields at 1.0 V/cm, no significant differences were seen between treatment and control groups in either TNT formation or cell growth (Fig 1D, E).

### Assessment of cell viability and DNA fragmentation following TTFields treatment

With TTFields application at 1 V/cm, a cytotoxic effect on cells was expected. However, as reported above, both MSTO-211H and VAMT continued to divide, even when monitored 24 hours following treatment. To confirm that the cells were indeed viable, we next performed cell viability assays at all timepoints on randomly selected fields of view using NucGreen Dead 480. In all cases, cell viability of both control and treatment groups was > 95% (Fig 1F,G), demonstrating no induction of cell death in the treated cells. Because TTFields exposure is known to affect cancer progression, we measured DNA fragmentation through the TUNEL assay. To confirm our earlier results with NucGreen Dead 480 and investigate cell viability at 150 kHz, we also repeated cell viability assays with NucGreen Dead 480 at both 150 kHz and 200 kHz at 1.0 V/cm.

Knowing that maximum TNT suppression occurred in MSTO-211H at 48 hours, we performed both assays at the 48 hour timepoint. For both TUNEL and NucGreen Dead assays, we noted minimal cell death with 2.4% and 1.8% mean cell death respectively at 150 kHz and 3.2% and 1% mean cell death at 200 kHz when referenced with a negative and positive control (Fig 1H, S2). As our findings of exponential cell growth and low cell death are in contrast to previous TTFields application studies, we repeated the experiments above at 1.0 V/cm and 200 kHz, but this time plated cells at a much lower density. In concurrence with others, we noted an 80% reduction in cell count in the TTFields-treated group when compared to control by the 72-96 hour timepoint (Fig S3, p=0.003)

### Effect of TTFields Exposure on actin polymerization and filament bundling

There are many unidentified molecular factors in the actin polymerization mechanism that form TNTs, including actin nucleators, elongators, bundlers, and destabilizers. In addition, there are membrane bound proteins involved in the process, and some of these components may differ between cell types. Filamentous actin forms the structural basis of the interior of TNTs. Because we observed a reduction in MSTO-211H TNTs with TTFields at 1.0 V/cm, and noting that tubulin depolymerization and polymerization has been observed to be directly impacted by TTFields treatment^15^, we next sought to determine what effects TTFields might have directly on actin at the polymer level. To accomplish this, we performed actin sedimentation experiments to examine both polymerization and bundling. Actin monomers in solution were combined with a KCl, MgCl_2_, and EGTA containing buffer to initiate polymerization, and for experimental samples, treated with TTFields at 1.0 V/cm-200 kHz with the inovitro device. After one hour of incubation, solutions were spun down and run on an SDS-PAGE. Surprisingly, there was no difference between samples treated with or without TTFields (Fig 2A,B). For both control and treated samples, actin was predominately found in the filamentous form. If TTFields did not directly alter actin polymerization, we considered a role for other components of the actin-based protrusion mechanism. As an initial experiment, we analyzed the actin bundling protein fascin to determine whether it was affected by TTFields. Again, there was no difference in the amount of actin bundling between TTFields-treated samples and controls, indicating that TTFields likely affect TNT formation by other factors in this system (Fig 2C,D).

**Figure 2.**
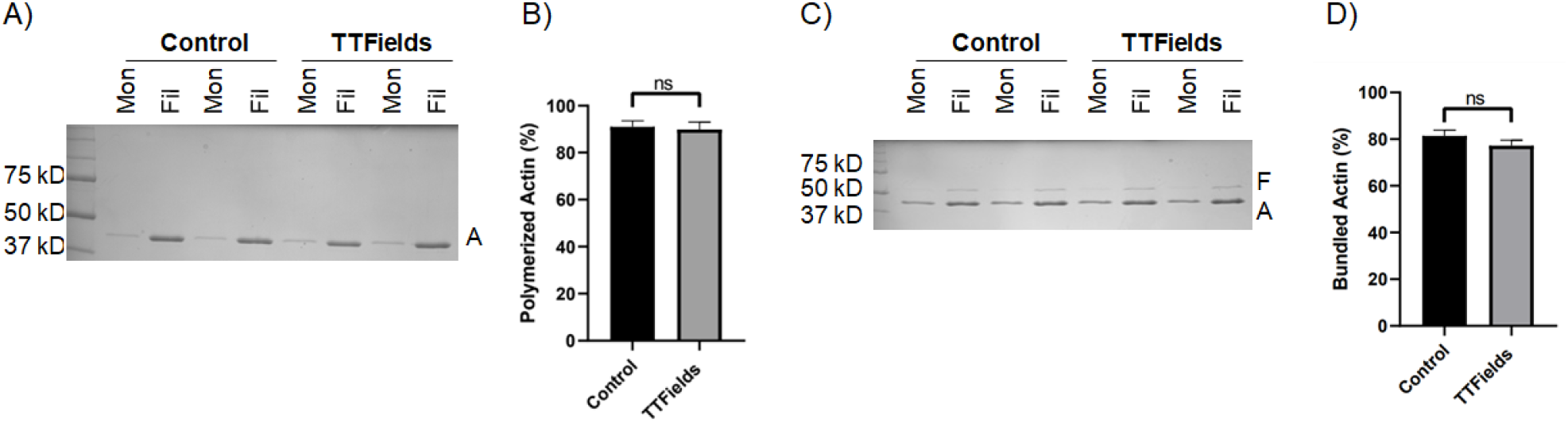
The effect of TTFields application on actin polymerization and actin filament bundling. (**A,B**) Sedimentation assays quantifying actin polymerization. Purified actin monomers were polymerized for 1 hour with TTFields (200 kHz, 1.0 V/cm, 37°C) and without TTFields (37°C) treatment. Reactions were centrifuged at 100,000 x g to pellet filamentous actin and analyzed by SDS-PAGE. Mon refers to monomeric actin (supernatant), Fil refers to filamentous actin (pellet). A indicates the actin protein band (42 kDa). (C,D) Co-sedimentation assays quantifying bundling of actin filaments by the bundling protein fascin. Pre-polymerized actin filaments were incubated with fascin for 1 hour with TTFields (200 kHz, 1.0 V/cm, 37°C) and without TTFields (37°C) treatment. Reactions were spun at low-speed (10,000 x g) to pellet bundles and analyzed by SDS-PAGE. The supernatant contains monomeric actin and individual filaments. The pellet contains bundled actin. F, A indicate fascin (55 kDa) and actin (42 kDa) protein bands. The gels (A,C) represent one representative experiment. The graphs (B,D) represent the average of three experiments, and the error bars are the standard deviation.

### The addition of chemotherapeutic agents to TTFields leads to reduced TNT formation and cell growth

TTFields are used clinically in patients concomitant with standard-of-care chemotherapy. The degree to which the interactions between and effects of TTFields and chemotherapy given together are synergistic has been shown when adding pemetrexed to cisplatin chemotherapy^22^. Demonstrating that TTFields exposure suppresses TNTs in MSTO-211H cells, we leveraged our ability to assess dynamic changes over time through continuous application of TTFields while capturing live-cell reaction during time-lapse microscopy. To do this, we utilized inovitro Live, a device that applies continuous TTFields while inserted into a tissue culture plate, and which is placed in an environmentally controlled microscope chamber. This experimental arrangement permits continuous viewing, imaging, and management of cells undergoing TTFields treatment in real time.

Thus, we posited that addition of standard-of-care chemotherapeutic drugs cisplatin (C) and pemetrexed (P) (Alimta) would work at least additively, and possibly synergistically, in combination with TTFields.

We performed a series of time-lapse experiments with 6 experimental groups: Control, TTFields only (1.0 V/cm, 200 kHz), Cisplatin w/o TTFields, Cisplatin+TTFields, Pemetrexed+Cisplatin w/o TTFields, and Pemetrexed+Cisplatin + TTFields (Fig 3A,B). When TTFields treatments were applied for 72 hours, a downward trend in TNT formation was observed compared to the control group (Fig 3A). This difference was most pronounced at the 48-hour timepoint, a result that was replicated from our original inovitro data in MSTO-211H (Fig 1A). Cell proliferation approximated an exponential growth curve for both the control and TTFields treatment groups, although the TTFields group had lower cell counts by 72-hours (Fig 3B). Next, we examined TNT formation and cell proliferation when the chemotherapeutic drug cisplatin was added at a physiologically relevant concentration (160 nM) to cells in culture. TNT formation was suppressed throughout the 72-hour period when compared to the control (Fig 3A). Cell growth was also suppressed, despite a higher cell count observed in the cisplatin group at the 0-hr timepoint (Fig 3B). When TTFields treatment at 1.0 V/cm and cisplatin were added concurrently, TNT suppression was even more pronounced, and this suppression again lasted for 72 hours. TNT formation was also suppressed in cells cultured with cisplatin and pemetrexed without TTFields at all timepoints when compared to the control (Fig 3A). Cell growth was similar to controls for 0 and 24 hours, but by 72 hours the cell growth of the cisplatin and pemetrexed treatment was suppressed (Fig 3B). When TTFields treatment was combined with cisplatin and pemetrexed, TNTs were also suppressed for the duration of the experiment similar to treatment with only cisplatin and pemetrexed. Cell growth under this condition was similar to controls at 0 and 24 hours, with cells in the treatment group ending at 72 hours with fewer cells than the control.

**Figure 3.**
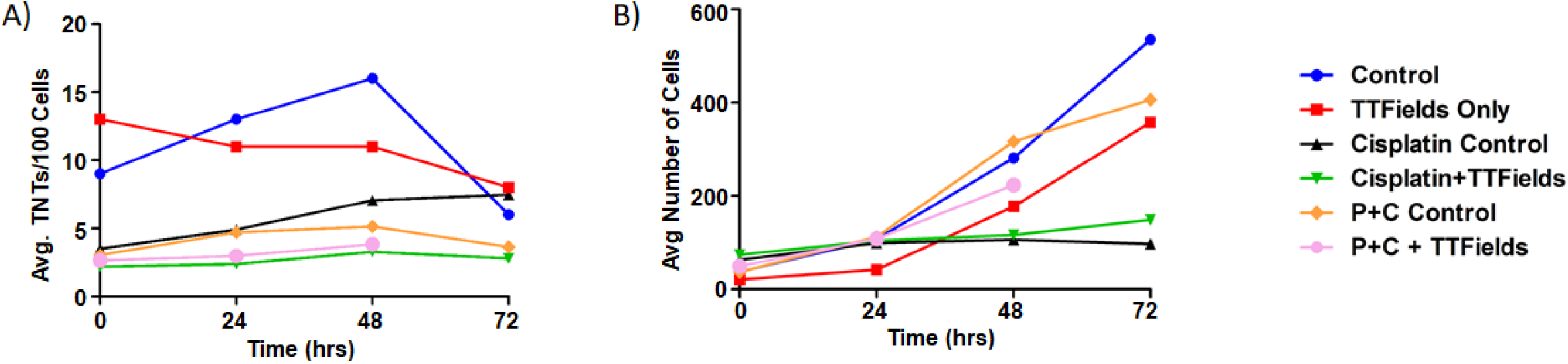
The effect of synergistic TTFields and chemotherapeutic exposure on MSTO-211H TNT formation and cell growth. A) TNT formation following treatment with cisplatin and cisplatin+pemetrexed over 72 hours. Intensity and frequency were set at 1.0 V/cm and 200 kHz respectively with bidirectional field delivery. B) Cell growth with chemotherapeutic reagents (C, cisplatin and P, pemetrexed) at 1.0 V/cm, 200 kHz, bidirectional. Results are indicative of one independent experiment (n=1) but with 45 technical replicates (TNTs/cell measured in multiple regions within the same experiment) averaged for each time period and condition.

### TNT Cargo Transport

TNTs allow for direct transfer of cell cargo and communication between cells. As TTFields applied at 1 V/cm suppressed formation of TNTs in MSTO-211H, we next sought to assess the effects of TTFields at these parameters on the ability of intact TNTs to mediate intercellular transport. We sought to track two kinds of TNT cargo: gondolas (bulges) representing cellular cargo being transported via TNTs that can be tracked with brightfield microscopy, and mitochondria, which we tracked using standard commercially available fluorescent labels. Gondolas were analyzed in MSTO-211H cells treated with no TTFields (control) and 200 kHz unidirectional, 200 kHz bidirectional, and 150 kHz bidirectional TTFields (Fig 4A). Images were captured every 60 seconds for one hour and analyzed by the Fiji-ImageJ Manual Tracking plugin. In the control group, the average velocity of TNT transport was 3.59 um/min. The average velocity of TNT transport was 3.94 um/min, 4.07 um/min, and 3.07 um/min for cells treated with TTFields delivered unidirectionally at 200 kHz, bidirectionally at 200 kHz, or bidirectionally at 150 kHz, respectively. These findings indicated that there were no observable differences in visible cargo velocities moving through TNTs in cells treated with or without TTFields.

**Figure 4.**
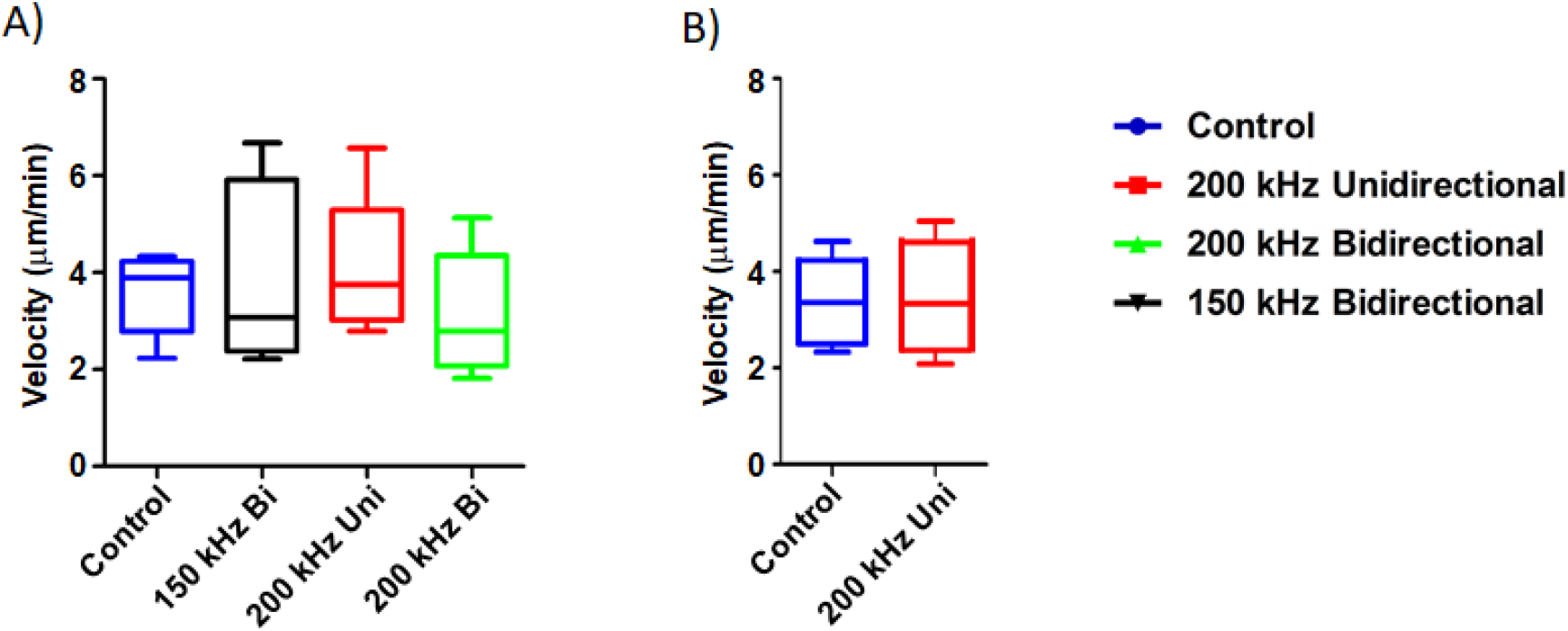
The effect of TTFields application on cargo transfer along TNTs. A) Cargo velocity with 1.0 V/cm, 150 or 200 kHz, unidirectional or bidirectional TTFields application. B) Mitochondrial velocity with 1.0 V/cm, 200 kHz unidirectional TTFields application. Results are indicative of three independent experiments (n=3).

Transport of mitochondria through TNTs has been extensively characterized to date^2^, and could indicate another way TTFields impact TNT functionality. MSTO-211H cells were stained with MitoTracker Orange (Thermo Fisher Scientific) and plated for optimal TNT formation. The following day they were either treated with or without TTFields applied unidirectionally at 1.0 V/cm and 200 kHz. Images were captured every 60 seconds for one hour, and fluorescently labeled mitochondria were analyzed using Fiji-ImageJ Manual Tracking plugin. In the control group, we showed that the average velocity of mitochondria was 3.32 um/min, with a standard deviation of 0.504 um/min, and with TTFields treatment an average velocity of 3.43 um/min, with a standard deviation of 0.17 um/min (Fig 4B). This finding indicated that similar to gondolas, there was no observable effect of TTFields on mitochondrial transfer in TNTs at the intensity and frequency that suppressed formation of TNTs.

### Spatial transcriptomic signatures of tumors treated with TTFields: Genetic effects of applying the TTFields to treat tumors in an *in vivo* animal model of mesothelioma

At present, there is no validated specific structural biomarker for TNTs, though there are proteins known to be upregulated in TNT formation in cancer phenotypes. Approaches to molecular analysis that could uncover TNT-specific biomarkers with high sensitivity would be an important advance for the field. At the same time, there are few studies reporting alterations in molecular pathways associated with TTFields-based treatment of cells or *in vivo* tumor models. We thus sought to leverage a spatial genomics approach to determine whether genes that have been associated with TNT formation and maintenance, are differentially expressed in a spatially distributed manner in intact tumors; and also to identify a convergent population of genes that are both differentially expressed following treatment using TTFields and also implicated in TNT biology. Within that context, to characterize alterations induced by TTFields at the genetic and molecular levels, and potential effects in particular on TNT-associated biomarkers, we performed spatial genomic analysis on an animal model of mesothelioma treated with TTFields, or alternately with heat as a sham for a negative control.

Eight total mice were injected with AB1 mesothelioma cells and assessed for tumor growth until they reached 200 mm^3^ in size. Four mice each were treated with TTFields using the inovivo™ device, or heat sham for a negative control, as described in Materials and Methods. Following TTFields or heat application, the mice were sacrificed, and the tumors were formalin fixed and paraffin embedded. One section from all eight tumors was adhered to a glass slide, as per Nanostring GeoMx instructions, from which a total of twelve regions of interest (ROI) were chosen. Six ROIs were from TTFields-treated tumors, and six from heat sham-treated tumors. These 6 ROIs were further divided into high or low Ki67 positive regions, as a measurement of mitotic index. NanoString’s GeoMx Digital Spatial Profiler system, with their mouse cancer transcriptome atlas (Catalog Number: GMX-RNA-NGS-CTA-4), was used to analyze the expression level of 1812 genes within our ROIs.

Analysis of gene expression showed that 22 of the CTA 1812 genes analyzed were differentially expressed (DEG) (Fig 5, Table 1). Broadly we found that the application of TTFields results in regulation of genes involved in cell adhesion and motility, PI3K-AKT signaling, and immune response; and to a lesser extent MAPK and MET signaling, and matrix remodeling-metastasis (Fig 5A). We focused on the subset of genes from the low Ki-67 ROIs, as their DEG was more pronounced. We reasoned that as the cells in these regions had a low rate of cell division, they were more affected by TTFields application, and thus potentially would be more likely to reveal genes that regulate TNT formation.

**Table 1.** DEG genes of TTFields treated tumors.

| Gene | Log2 Fold Change | p-Adjusted |
| --- | --- | --- |
| LYZ | 1.9942 | 2.20E-07 |
| COL1A1 | 1.9875 | 6.49E-05 |
| HLA-DRB | 1.9839 | 1.18E-03 |
| COL1A2 | 1.5858 | 5.55E-08 |
| CSF1 | 1.1208 | 8.69E-08 |
| COL3A1 | 1.1203 | 2.73E-02 |
| ACTA2 | 1.0931 | 4.45E-12 |
| CX3CR1 | 1.0117 | 4.60E-12 |
| C1R | 1.0034 | 8.69E-08 |
| HDAC5 | -1.0960 | 4.20E-04 |
| PKM | -1.1221 | 2.74E-02 |
| INHBA | -1.1502 | 1.01E-11 |
| VEGFA | -1.1652 | 7.21E-05 |
| CD9 | -1.1966 | 7.56E-03 |
| MASP1 | -1.3157 | 3.36E-04 |
| SLC2A1 | -1.3392 | 3.05E-03 |
| STC1 | -1.4611 | 3.62E-03 |
| CEBPB | -1.4803 | 7.25E-05 |
| LRP1 | -1.5079 | 2.35E-03 |
| HGF | -2.0459 | 2.52E-06 |
| FST | -2.2774 | 6.84E-06 |
| TNC | -2.4863 | 2.00E-06 |

**Figure 5.**
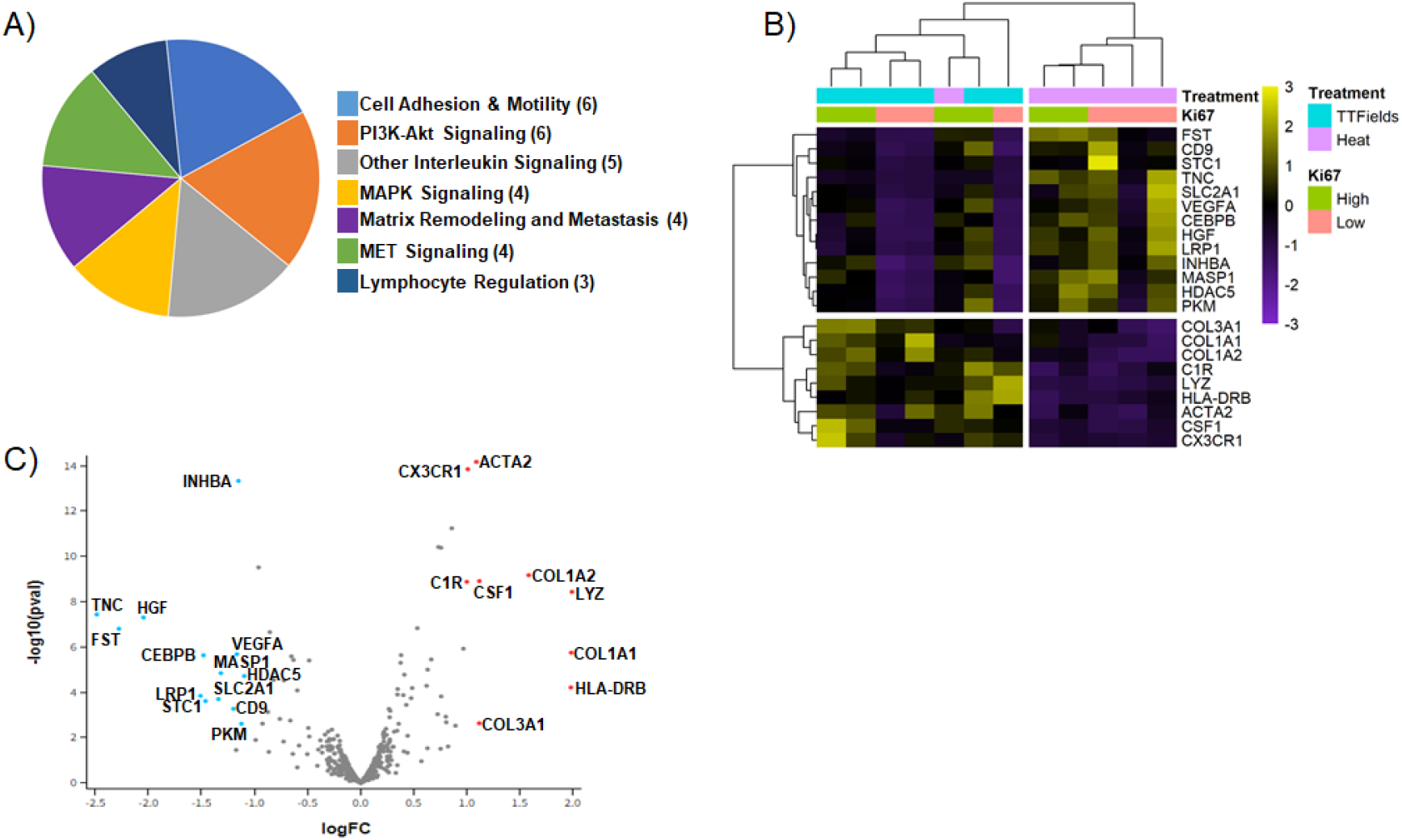
DEG genes of TTFields treated tumors. (A) Categories of genes found to be differentially expressed. () indicates the number of genes, that fall into a given category. (B-C) Heatmap and Volcano plot generated by spatial omics analysis.

The genes most prominently affected (downregulated) in TTFields treated tumors as compared to the heat controls, were TNC, FST, and HGF (Fig 5 B, C, Table 1). TNC is aglycoprotein involved in the epithelial-to-mesenchymal transition, and was previously found by our group to be regulated in TNT promoting conditions^23^. In contrast, upregulation was most prominent for HLA-DRB, LYZ, COL1A1, and COL1A2. LYZ is associated with neutrophil degranulation and host defense peptides, whereas COL1A1, COL1A2 along with HGF, TNC, and VEGFA are all part of the PI3K pathway, which plays an important role in cancer progression, and has been implicated in TNT regulation^24^. Expression of immunogenic markers with implications for efficacy of immune-oncology therapeutic strategies were also found, and included CX3CR1, which was upregulated in TTFields-treated tumors overall, as well as the aforementioned HLA-DRB, C1R and COL3A1. Markers of angiogenic activity, such as VEGFA, which are also implicated in EMT, hypoxia signaling pathways and cell adhesion and motility, also were notably downregulated. In sum, application of TTFields altered a spectrum of metabolic and molecular signaling pathways that are well established in cell proliferation and division, ancillary pathways associated with construction and maintenance of the tumor matrix, while at the same time upregulating certain immunogenic markers.

## Discussion

TTFields are low intensity (1-3 V/cm) alternating electric fields applied at frequencies ranging from 100-400 kHz^21, 22^ and have been shown to impact polar proteins during cellular replication, specifically tubulin. Because the main component of TNTs is F-actin molecules, which have an electrochemically polar nature, we decided to study the effect of TTFields treatment on TNT formation in mesothelioma. In this study, we investigated the ability of TTFields to affect TNT formation and function in MPM, and also evaluated genetic signatures affected by TTFields treatment of MPM in an animal model. We found that TTFields significantly suppressed formation of TNTs in the biphasic (epithelioid plus sarcomatoid) form of MPM represented by the MSTO-211H cell line, when TTFields were applied at standard intensity of 1.0 V/cm, 48 hours after initiation of treatment. No significant differences were seen at 24 hours, nor subsequent to 48 hours, when cell crowding under cell culture conditions naturally leads to fewer TNTs. We found no detectable effect on TNTs with the pure sarcomatoid cell line VAMT. We assessed free actin, in monomeric and filamentous form, and found no detectable differences due to TTFields in this context. Spatial genomic assessment of intact MPM tumors following TTFields detected a notable upregulation of immuno-oncologic biomarkers, with concurrent downregulation of multiple metabolic, cell signaling, and cell growth pathways associated with dysregulation of MPM and other cancers. Some of these signals have also been implicated by our team and others in TNT activity in MPM and similar cell types.

TNTs are F-actin based protrusions, involving dynamic microtubules, either in active or passive fashion, in trafficking of cytosolic cell contents from cell-to-cell. TNTs have been shown to be disrupted primarily through knockdown or inhibition of protein complexes that promote actin formation, such as M-sec (TNFaip2), Arp 2/3, and others^25, 26, 27, 28^. The application of TTFields has already been shown to disrupt mitosis in actively dividing cells via their effect on microtubules, using a non-pharmacologic approach. Because TNTs are composed of polar actin subunits, a significant disruption of TNTs would suggest that actin is required for nanotube stability. G-actin subunits, the main component of TNTs, contain a distinct polarity. TTFields could be used to force G-actin subunits to align along the electric field instead of polymerizing. However, as there was no effect of TTFields on cell-free forms of actin, our findings suggested a more selective mechanism of TTFields. Indeed, when controlling for other parameters, the maximum suppression of TNT occurring unidirectionally vs bidirectionally may indicate that the orientation of the affected TNT component as well as its identity may play important roles in TNT formation. Thus, further studies unifying the mechanism of TTFields and the ultra-structure of TNTs are needed. Overall, we have demonstrated that TNTs are more likely to be affected because of either microtubule disruption or other associated cell machinery than actin.

Preclinical models of TTFields have demonstrated their ability to induce cell death over time. In the current *in vitro* study, overall cell viability remained above 95% in both the control and treatment groups at both intensities and at all timepoints when 40,000 MSTO-211H cells were seeded one day before treatment started.

However, when MSTO-211H were seeded at a lower density and exposed to TTFields, markedly reduced cell counts were observed at 72-96 hours of exposure (Fig S3), indicating that TTFields cytotoxic effect, at least in vitro, is affected by cell density. Ultimately, TTFields should be used in conjunction with other forms of cancer therapy, such as radiation therapy or chemotherapy to achieve maximum efficacy.

MPM is an ideal model for *in vitro* study and characterization of TNTs. It thus proved especially useful here, with additional value and background that MPM cells lines, including MSTO-211H had previously been evaluated after exposure to TTFields. Giladi et al. conducted a study on the optimal inhibitory frequencies and intensities of various cell lines exposed to TTFields. It was found that of the 30 cell lines tested, MSTO-211H was categorized as sensitive to the cytotoxic effects of TTFields^15^. Such a finding could explain the discrepancy in TNT formation between MSTO-211H and VAMT, suggesting that properties specific to individual cell lines could allow for resistance to TTFields treatment. While optimal inhibitory frequency/intensity of VAMT sarcomatoid MPM was not measured in the study, its behavior under TTFields, specifically a non-significant difference in TNT formation, suggested that it is less sensitive relative to MSTO-211H.

The precise cellular mechanism(s) and identity of molecular machinery complex(es) necessary for TNT-mediated intercellular trafficking have not yet been identified. It is conceivable that the process mirrors the ones seen in other filamentous membrane-based protrusions, and it is equally conceivable that the process of TNTs may be cell type-dependent. Is actin necessary for the function of TNTs, or just the structure? Does actin polymerization correlate with TNT stability, in addition to function? Answers to these questions could provide insight into specific molecular markers involved in TNT formation as well as targeted therapy options in clinical practice. One idea stems from the role of the Arp 2/3 complex, which serves as a nucleation site for actin filaments by binding to the side of one filament and subsequently acting as a template for another filament, which is added at a 70 degree angle relative to the first filament^29^. While Arp 2/3 has not directly been studied, the Rho GTPase protein family has been observed to localize multiple proteins, including Arp 2/3, that can then serve as potential nucleation sites for actin filaments^30^. Indeed, TTFields application has been shown to activate the Rho-ROCK pathway and promote reorganization of the actin cytoskeleton, which may explain our findings on TNT suppression in MSTO-211H^31^.

The use of TTFields performed *in vitro* may provide insight into TNT biology. However, we sought to move a step beyond that by leveraging an even newer version of the technology that permits tailored treatment of TTFields *in vivo* to tumors in animal models. We utilized this technology to accurately treat multiple MPM tumors, then further leveraged a spatial genomics approach to uncover the spatial geography of TTFields effect, and determine what links would exist, if any, between differential expressed genes and our current and past findings of TNTs *in vitro*. The findings from spatial genomic analysis overall were highly notable for uncovering classes of immuno-oncologic response genes that were upregulated following TTFields exposure, in comparison to heat sham-treated tumors. The clinical implication for this finding is important because it is not yet established which set of cancer-directed therapies match best with TTFields, and in what sequence (prior to, during, or following each other), to produce best clinical response. Upregulation of factors such as CSF1 (macrophage colony stimulating factor-1, a cytokine responsible for macrophage production and immunoresponse), CX3CR1 (chemokine signaling), and HLA-DRB (lymphocyte trafficking and T cell receptor signaling), with concurrent modulation of the tumor microenvironment (TME) mediated by increased expression of collagens COL1A1, COL1A2, and COL3A1, may induce an inflammatory niche susceptible to cutting edge therapeutic including immune checkpoint inhibitors. These results suggested a possible role of combining TTFields with immunotherapy in creating a more drug-targeted friendly TME.

Beyond those results, it is the downregulated set of genes that is most prominent in identifying signals that could explain why formation of TNTs in biphasic MPM was suppressed by TTFields. Numerous classes and specific genes involved in cell adhesion and motility or in epithelial-to-mesenchymal transition (EMT) were downregulated by TTFields-treated MPM, including most prominently Tenascin C (TNC) and vascular endothelial growth factor A (VEGFA). We have previously reported that Tenascin C, a modulator of cell invasive potential, is upregulated in mesothelioma cells primed in cell culture conditions conducive to TNT formation^23^. Furthermore, transition of mesothelioma cells to EMT is strongly associated with a sharp rise in TNT formation. We have also reported on the intercellular transport of VEGF, a finding that implicates TNTs in other cancer-provoking processes including angiogenesis. In regards to the Arp2/3 complex, none of these genes were included in the transcriptome atlas we used for this study. RhoA and B expression were assayed, but no differential gene expression was observed. The data signals shown using spatiotemporal analysis produced an overview of a TME that was clearly reconfigured by TTFields treatment, one that has crossover with factors associated with TNT formation and function as shown *in vitro*. Future studies will unravel the role of individual factors or groups involved in TNT formation and maintenance, and prove whether they are necessary and sufficient for these processes.

Limitations of this study include uncertainty of factors that are necessary, sufficient, and crucial to formation and maintenance of TNTs both *in vitro* and *in vivo*. In this context, it is uncertain as of yet why TNTs in the biphasic (epithelioid and sarcomatoid) MSTO-211H cell line responded effectively to TTFields treatment, but TNTs in the purely sarcomatoid cell line VAMT did not. All inovitro experiments were limited by the maximum size of the 22 mm coverslip used to culture cells for TTFields treatment; and only at this diameter could the coverslip fit into the ceramic dish for TTFields delivery. Thus, a delicate balance existed between plating too high a density of cells approaching confluency versus plating too few of cells such that growth rate was suboptimal.

In this study, we report novel cellular and molecular effects of TTFields in relation to tumor communication networks enabled by TNTs and related molecular pathways. TTFields significantly suppressed formation of TNTs in biphasic malignant mesothelioma (MSTO-211H). Spatial genomic assessment of TTFields treatment of intact mesothelioma tumors from an animal model shed new light on gene expression alterations at the transcriptomic level that imply how TTFields may provide synergy with chemotherapy and immunotherapeutic strategies. These results position TNTs as potential therapeutic targets of TTFields and also identify the use of TTFields to remodulate the tumor microenvironment and enable a greater response to immunotherapeutic drugs.

## Materials and Methods

### Cell Lines and Culture

MSTO-211H cells are a biphasic MPM cell line that was purchased from the American Type Culture Collection (ATCC, Rockville, MD, USA) for use in this study. VAMT is a sarcomatoid MPM cell line that was authenticated prior to use. Both cell lines were grown in RPMI-1640, supplemented with 10% Fetal Bovine Serum (FBS), 1% Penicillin-Streptomycin, 1x GlutaMAX (all from Gibco Life Technologies, Gaithersburg, MD, USA), and 0.1% Normocin anti-mycoplasma reagent (Invivogen, San Diego, CA, USA). Cells were negative for mycoplasma infection, and were maintained in a humidified incubator at 37°C with 5% carbon dioxide. Cell viability was assayed by treating cells with NucGreen Dead 488 ReadyProbes Reagent (Invitrogen, Carlsbad, CA, USA), imaging seven random fields of view, and quantifying these fields. Apoptosis and DNA fragmentation were assayed with Click-iT TUNEL Alexa Fluor 488 Imaging Assay (Thermo Fisher Scientific, Waltham, MA, USA) according to the manufacturer’s instructions.

### inovitro TTFields Treatment

An inovitro™ device, provided by Novocure, Ltd (Haifa, Israel), was used to apply continuous bidirectional TTFields treatment to cells. One day prior to treatment with TTFields, 22-mm plastic cell-culture treated coverslips (Thermo Fisher Scientific Nunc Thermanox, Waltham, MA, USA) were placed inside sterile ceramic dishes. MSTO-211H cells (40,000) in 2 ml of growth media were plated onto the coverslips, and the dishes were placed in a base plate in a humidified incubator at 37°C with 5% carbon dioxide overnight. To apply TTFields to the cells, the ceramic dishes were connected to an inovitro Generator Box. inovitro software controls and monitors the electrical resistance, voltage, and current in real time, while the temperature in the incubator is directly correlated with the intensity of the electric field. The temperature was set at 32°C to deliver an intensity of 0.5 V/cm and at 26.5°C for an intensity of 1.0 V/cm^20^. Additionally, the frequency of the electric field was set at 200 kHz for all conditions in both cell lines, barring any initial frequency testing and cell viability assessment. All intensity values were expressed in root mean square (RMS) values to illustrate the conventional depiction of alternating current measurements in physics fields. The treated group was exposed to TTFields for 72 hours in both 0.5 V/cm and 1.0 V/cm experiments. For the 1.0 V/cm experiments, the TTFields were shut off at 72 hours, and the cells were incubated for another 24 hours to assess recovery of TNTs. Cells in the control group were not treated with TTFields, but were plated as described above and placed in an incubator at 37°C with 5% carbon dioxide for the duration of the experiment. The low density experiments were run as above with the exception that only 10,000 cells were plated onto a coverslip, and TTFields application followed 3 hours later.

### TNT Analysis and Quantification

Quantification and visual identification of TNTs were performed as described previously^2, 7, 19, 23, 32^. Briefly, these parameters included (i) lack of adherence to the substratum of tissue culture plates, including visualization of TNTs passing over adherent cells; (ii) TNTs connecting two cells or if extending from one cell were counted if the width of the extension was estimated to be <1000 nm; and (iii) detection of a narrow base at the site of extrusion from the plasma membrane. Cellular extensions that were not clearly identified with the above parameters were excluded. Still images and time-lapse videos were analyzed using Fiji-ImageJ software. The Fiji-ImageJ Multi-point tool was used to quantify TNTs and cell number following the criteria detailed above; and the TNT index was calculated as the number of TNTs per 100 cells. The X, Y coordinate function was used to calculate the length of TNTs, using a conversion of 0.335 μm/pixel with a 20x objective.

### Time-lapse Microscopic Imaging with Concurrent Continuous Administration of TTFields using inovitro Live

An inovitro Live™ device, provided by Novocure, Ltd (Haifa Israel), was used to apply continuous unidirectional or bidirectional TTFields exposure to cells. One day prior to treatment, 40,000 MSTO-211H cells were plated onto a 35 mm high wall, glass bottom dish (Ibidi, Gräfelfing, Germany), and allowed to adhere overnight. For the unidirectional and bidirectional experiments, the glass bottom dish was coated with Poly-D-Lysine (Millipore Sigma, Burlington, MA) at a concentration of 1mg/µm for 1 hour then dried for 2 hours prior to plating. The next day, an inovitro Live insert was positioned in the 35 mm dish, and placed in the microscope chamber. The plate was connected to an inovitro Live cable, and a heating element was added on top of the dish cover to minimize condensation from heat generated by TTFields. The cable was then connected to an inovitro Live Generator, and the software controlled the delivery of an electric field in either one (unidirectional) or two (bidirectional) directions at an intensity of 1.0 V/cm and either 150 or 200 kHz. Media was changed every 24 hours, during which TTFields were paused and then resumed once the cells were placed back into the incubator. The cells for the control group were plated as described above and placed in the microscope chamber at 37°C, without TTFields, for the duration of the experiment. Seven Fields of View (FOV) were selected every 24 hours, up to 72 hours and both cell proliferation and TNT formation were quantified.

As an additional experimental arm, MSTO-211H cells were also treated with cisplatin (160 nM) and pemetrexed (24 nM) in conjunction with TTFields application using pre-treated ibidi plates. During these experiments, images were acquired for 4 hours at 2min/frame, and this process repeated every 24 hours, up to 72 hours total. Both cell proliferation and TNT formation were subsequently quantified as described above.

Still images and time-lapse videos were taken on a Zeiss AxioObserver M1 Microscope. In order to deliver TTFields at an intensity of 1.0 V/cm, the microscope chamber temperature was set to 26.5 °C. Images were taken on a 20X PlanApo-Chromat objective with a numerical aperture of 0.8. We used a Zeiss Axio Cam MR camera with 6.7×6.7 µm width, and spatial resolution (dx=dy) at 20X was 0.335 µm/pixel. Images were acquired on Zen Pro 2012 software in brightfield.

### Cargo and Mitochondria Transfer

Cargo Transfer within TNTs was calculated using the Manual Tracking Plugin on Fiji-ImageJ. The X, Y coordinate of each cargo was recorded over time, and exported to a spreadsheet. To calculate velocity of cargo, X and Y pixel measurements were converted into microns using the scale factor 0.335 µm/pixel (20x objective). Then, the distance formula was implemented for Xn and Yn values, where n is any subsequent location of the cargo in relation to the first location, X1 and Y1. This process was repeated for each cargo track to calculate distance. Finally, each distance was divided by the time interval between frames. To track mitochondria, MSTO-211H cells were stained with MitoTracker Orange CMTMRos (Thermo Fisher Scientific, Waltham, MA, USA) and followed the same experimental setup and analysis as described above.

### Actin and Fascin Purification

Actin was purified from chicken skeletal muscle by one cycle of polymerization and depolymerization using standard protocols in the field (Spudich et al.). It was then filtered on Sephacryl S-300 resin (GE Healthcare) in G-buffer (2 mM Tris (pH 8.0), 0.2 mM ATP, 0.5 mM DTT, 0.1 mM CaCl_2_) to obtain actin monomers, and stored at 4°C. Human fascin-1 was expressed with an N-terminal glutathione s-transferase (GST) tag and a TEV cleavage recognition sequence from the pGV67 plasmid in BL21 DE3pLysS competent cells. Transformants were grown in 1 L of LB broth, induced at OD_600_ ∼0.6 with 0.5 mM IPTG, and shaken overnight (200 rpm, 17°C). To purify fascin, cell pellets were resuspended in lysis buffer (50 mM Tris, pH 8.0, 500 mM NaCl, 1 mM DTT) and sonicated. Lysed cells were centrifuged (∼30,000 x g, 4°C) for 40 minutes to isolate the soluble cell components. Samples were rotated with glutathione agarose resin (pH 8.0) for 1 hour at 4°C, washed, and eluted (50 mM Tris, pH 8.0, 100 mM NaCl, 1mM DTT, 100 mM glutathione). Eluted fractions were incubated with TEV protease (1.6 µM) for GST tag cleavage and dialyzed into glutathione-free buffer overnight. To remove GST contaminants and TEV protease, samples were filtered through glutathione resin followed by amylose resin. Collected flow throughs were concentrated using centrifugal filters (MilliporeSigma Amicon, MWCO 30K). Samples were frozen in liquid nitrogen and stored at -80°C.

### Actin Polymerization and Bundling Sedimentation Assays

Actin was polymerized at 37°C in KMEI buffer (50 mM KCl, 1 mM MgCl_2_, 1 mM EGTA, 10 mM Imidazole pH 7.0) for 1 hour with and without 1.0 V/cm inovitro device TTFields treatment. Samples were centrifuged at 100,000 x g for 30 minutes at 4°C to separate filaments and monomers. Supernatant and pellet fractions were analyzed via SDS-PAGE (12% acrylamide). Gels were then stained with Coomassie Blue for 1 hour and destained for at least 6 hours (10% ethanol, 7.5% acetic acid). Band intensities were quantified via densitometry using Fiji-ImageJ. For bundling, actin (15 µM) was first polymerized for 1 hour at 37°C in KMEI buffer. The assembled filaments were diluted to 3 µM and added to a solution with fascin (300 nM). After 1 hour with and without 1.0 V/cm TTFields treatment, samples were centrifuged at 10,000 x g for 30 minutes at 4°C to pellet bundled actin. SDS-PAGE and band quantification were carried out as described previously.

### Spatial Genomics

Blocks of formalin-fixed paraffin-embedded (FFPE) mesothelioma tumors that were treated with sham heat or TTFields were generously provided by Novocure, LLC for Nanostring GeoMx spatial transcriptomic analysis. In brief, eight female mice (Mus Musculus species, strain C57BL, aged 13 weeks) were subcutaneously injected with AB1 mouse mesothelioma cells. After tumors formed, mice were treated with heat or TTFields using the inovivo device (Novocure, Ltd) for a total of 14 days: 7 days of treatment, 2 days of rest, and 7 days of additional treatment. The tumors were excised, formalin fixed and paraffin embedded, and sent to our lab. With these tumor blocks, one 5 µm section from each tumor was placed on a glass slide for Nanostring GeoMx analysis (Seattle, WA). The slide was incubated with Ki-67 antibodies and the GeoMx Mouse Cancer Transcriptome Atlas panel of 1,812 RNA probes. Regions of interest (ROIs) were chosen, and the unique DNA indexing-oligonucleotide tags were cleaved from the RNA probes within the ROIs. These tags were then sequenced and analyzed with GeoMx DSP software.

### Statistical Analysis

#### inovitro Experiments

Due to lower sample sizes and skewed distributions of TNTs/cell, heteroscedastic t-tests were performed to assess significance in differences between TNTs/cell. Significance tests were performed on GraphPad Prism (GraphPad Software, Inc., La Jolla, CA, USA). P-values less than 0.05 indicated statistically significant differences; and error bars were included in graphs to depict standard error.

#### Bidirectional versus Unidirectional inovitro Experiments

The number of TNTs/cell after TTFields exposure was compared within treatment groups as a function of time using a linear mixed model to account for the repeated measures at each timepoint and treatment condition within each experiment. A compound symmetry correlation structure was assumed. Least squares means and standard errors are reported. Overall tests and pairwise comparisons are reported; and no adjustments for multiple comparisons were made. Data were analyzed using SAS 9.4 (Cary, NC) and p-values <0.05 were considered statistically significant.

#### Spatial Genomics

A Wald test was performed to assess significance in differentially expressed genes from TTFields vs heat treated mice using the Deseq package in R (R Foundation for Statistical Computing, Vienna, Australia). For each p value generated, a Benjamini-Hochberg adjusted p-value was acquired to reduce false-positive rate and reported.

#### Animal Use and Ethical Approval

This study was performed in strict accordance with the recommendations in the Guide for the Care and Use of Laboratory Animals of the National Institutes of Health. All of the animals were handled according to approved institutional animal care and use committee (IACUC) protocols (QSF-GLP-059) of Novocure. The protocol was approved by the Israeli National Committee Council for Experiments on Animal Subjects (IL-19-12-484). All surgery was performed under ketamine-xylazine anesthesia, and every effort was made to minimize suffering.

Animals specifically used were of the Mus Musculus species (strain C57BL), female at 13 weeks, with no genetic modification, supplied by Envigo (Jerusalem, Israel, catalog number 2BALB/C26).

#### Adherence to community standards

ARRIVE and ICJME guidelines were followed for this work.

## Acknowledgements

Research reported in this publication was supported by the National Center for Advancing Translational Sciences of the National Institutes of Health Award Number UL1-TR002494. The content is solely the responsibility of the authors and does not necessarily represent the official views of the National Institutes of Health. We thank the American Association for Cancer Research (AACR) for funding support through the AACR-Novocure Tumor-Treating Fields Research Award (grant number 1-60-62-LOU), as well as the Masonic Cancer Center (MCC) through the MCC spatial grant. Additional sponsors of research work in this field in the Lou Lab include: the Minnesota Ovarian Cancer Alliance (MOCA); The Randy Shaver Cancer Research and Community Fund; the Litman Family Fund for Cancer Research; the Mu Sigma Chapter of the Phi Gamma Delta Fraternity, University of Minnesota. We would like to thank the University of Minnesota Genomics Center and the University of Minnesota University Imaging Center (UIC) for their resources and technical support. We thank Antonia Martinez-Conde, Boris Brant, Adi Haber, Eyal Dor-On, Yaara Porat, and Moshe Giladi from Novocure, Ltd for technical assistance including with time lapse experiments analyzed by our team, and by providing inovivo-treated tumor specimens for spatial transcriptomic analysis. We thank Fernanda Rodriguez and Grant Barthel for their technical and logistical support regarding spatial genomics. We thank Michael Franklin, MS from the Division of Hematology, Oncology and Transplantation at the University of Minnesota for helpful suggestions and assistance in editing this manuscript. The authors apologize that not all pertinent and seminal references in the field could be discussed or cited in this article due to space limitations.

## Financial Disclosures

E.L. discloses research collaborations and consulting for Novocure, LLC in 2018-21, and grant funding via the AACR-Novocure Tumor-Treating Fields Research Grant 2019-2022; reimbursement for travel from GlaxoSmithKline, Inc. for giving a research presentation in 2016; consultation fees for Boston Scientific in 2019; unpaid consultation for Nomocan Pharmaceuticals; and membership on the Scientific Advisory Board for Minnetronix, LLC (unpaid). This work was done in collaboration with team members from Novocure, LLC.

**Supplemental Figure 1:**
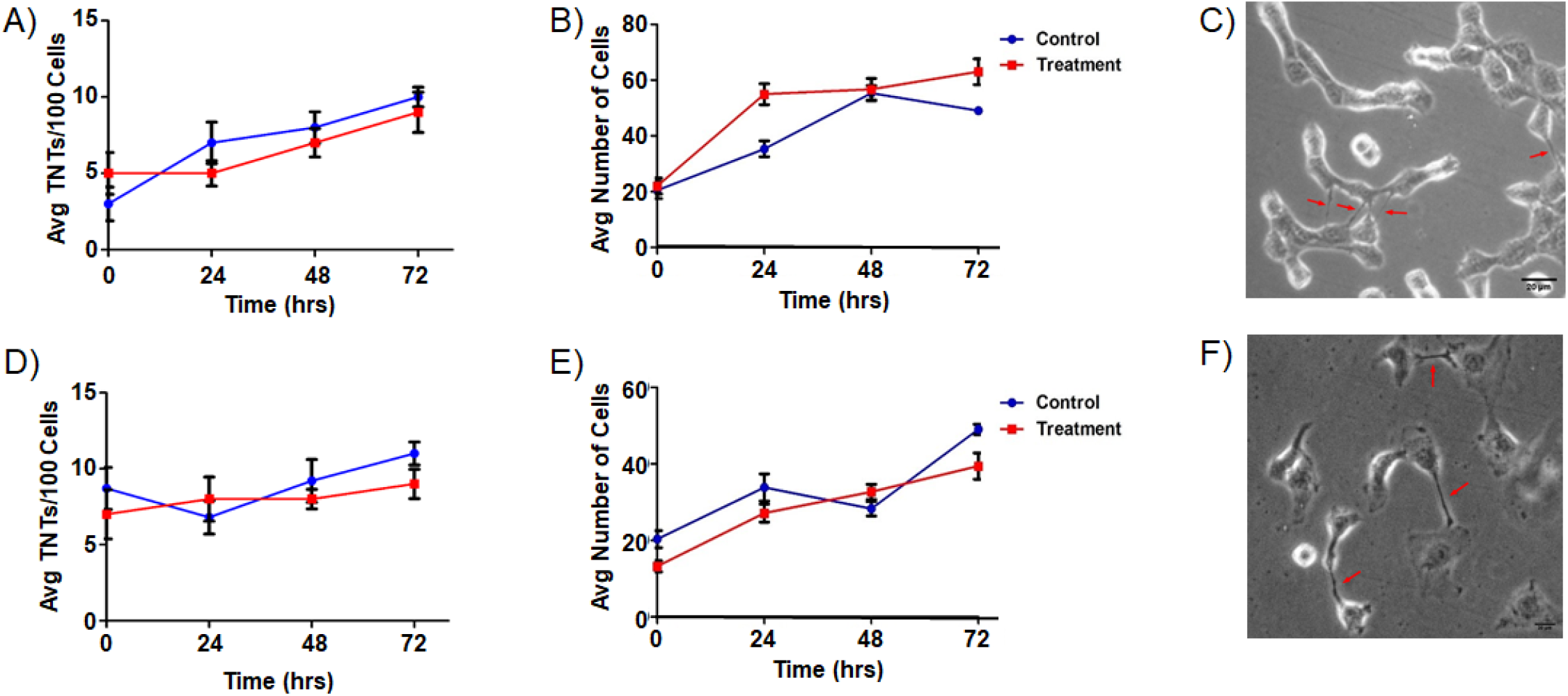
TTFields delivered at low intensity (0.5 V/cm) have no effect on TNT formation or on cell proliferation in MSTO-211H or VAMT mesothelioma cells. TNT formation and (B) cell growth in MSTO-211H with TTFields delivered at 0.5 V/cm, 200 kHz. (C) MSTO cells. Arrows point to TNTs. (D) TNT formation and (E) cell growth in VAMT with TTFields delivered at 0.5 V/cm, 200 kHz. (F) VAMT cells. Arrows point to TNTs.

**Supplemental Figure 2:**
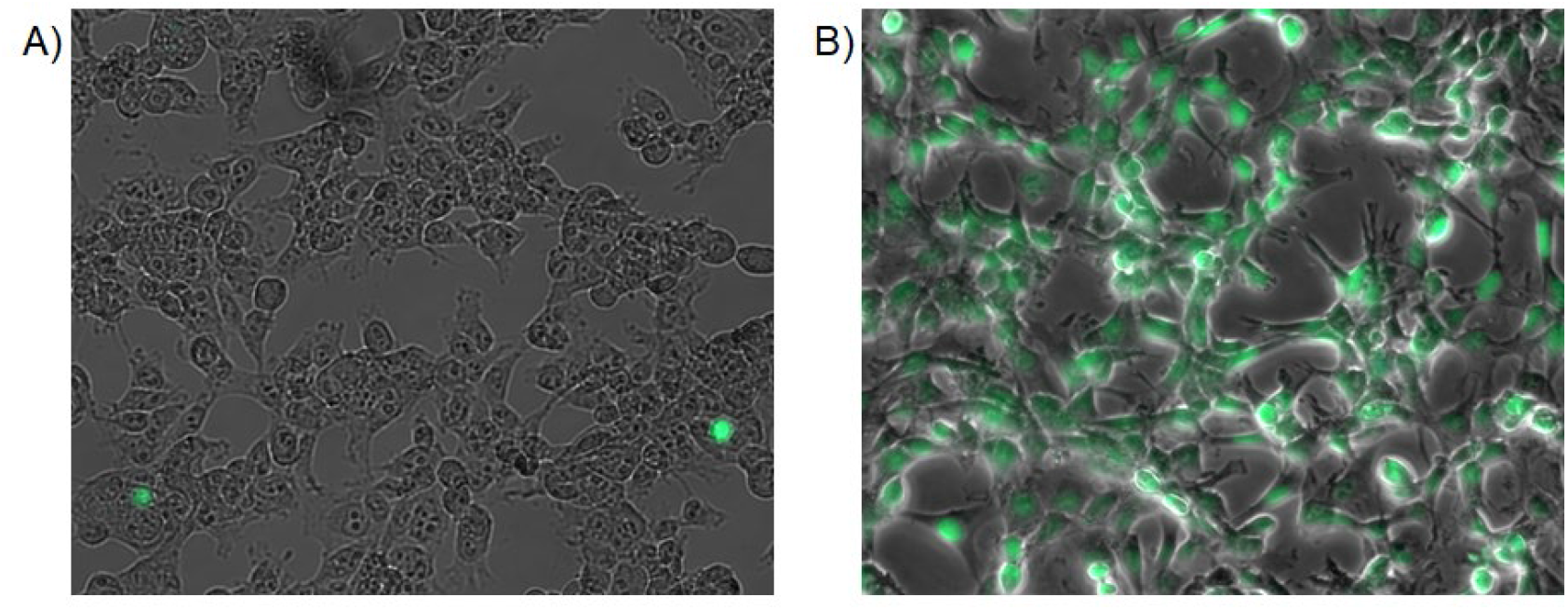
Representative images of the TUNEL assay in MSTO-211H. A) TUNEL assay in MSTO-211H after 48 hours of TTFields application or B) DNaseI treated positive control. Images were taken on a Zeiss AxioObserver M1 at 20X, with spatial resolution (dx=dy) at 0.335 um/pixel. Images were acquired on Zen Pro 2012 software.

**Supplemental Figure 3:**
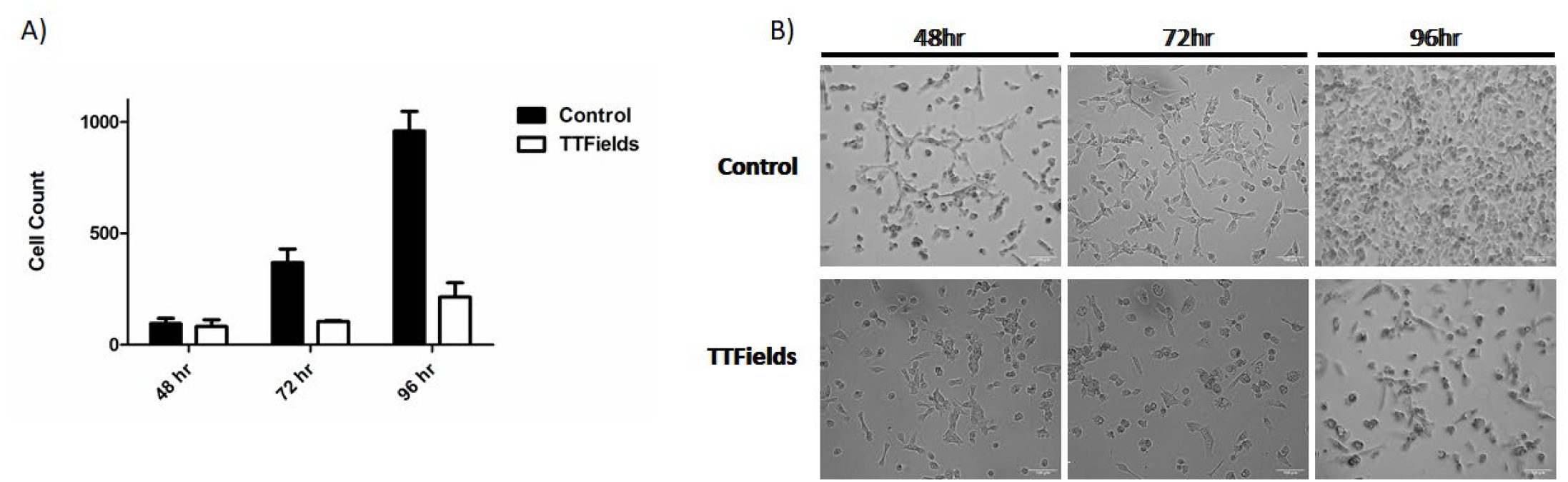
Cell count in MSTO-211H at 1 V/cm, 200 kHz, and seeded at 10,000 cells. (A) MSTO-211H cells were seeded at 10,000 cells/ml and treated with TTFields with the above specified parameters for 96 hours. Cell count was measured every 24 hours, starting at the 48 hour timepoint (n=3).Significance was assessed with heteroscedastic t-tests with three independent experiments performed, p=0.003 at 96 hours and p=0.048 at 72 hours B) Representative images of MSTO-211H cells at the 48, 72, 96 hour timepoints.

